# Redoxyme: a lightweight graphical user interface for standardized calculation of antioxidant enzyme activities

**DOI:** 10.64898/2026.02.05.703993

**Authors:** Geovanna Carvalho de Freitas Soares, Anna Lidia Nunes Varella, Heberty Tarso Facundo

## Abstract

Oxidative stress results from excessive accumulation of reactive oxygen species (ROS) and plays a central role in numerous physiological and pathological processes. Accurate quantification of antioxidant enzyme activities is therefore essential in redox biology research. However, data analysis for commonly used assays, such as superoxide dismutase (SOD), catalase (CAT), and glutathione peroxidase (GPx), is frequently performed using spreadsheets or manual calculations, which are time-consuming and prone to error. Here, we present Redoxyme, a free, open-source, Python-based graphical user interface designed to standardize and automate the calculation of antioxidant enzyme activities. The software integrates protein normalization, enzyme-specific calculation routines, data visualization, and Excel export within an intuitive interface that does not require programming expertise. Redoxyme was validated using experimental data obtained from animal tissues (rats and mice), demonstrating excellent agreement with manual calculations and established analytical methods. Redoxyme provides a practical solution for improving reproducibility and efficiency in antioxidant enzyme activity analysis. The software is currently distributed as a standalone executable for Windows (locally installed), and an interactive web-based calculator implemented in Streamlit, enabling direct use without local installation. The source code and version-controlled development history are openly accessible via GitHub, promoting transparency, reproducibility, community-driven improvements, and can, in principle, be adapted for other operating systems.

**Graphical Abstract:** 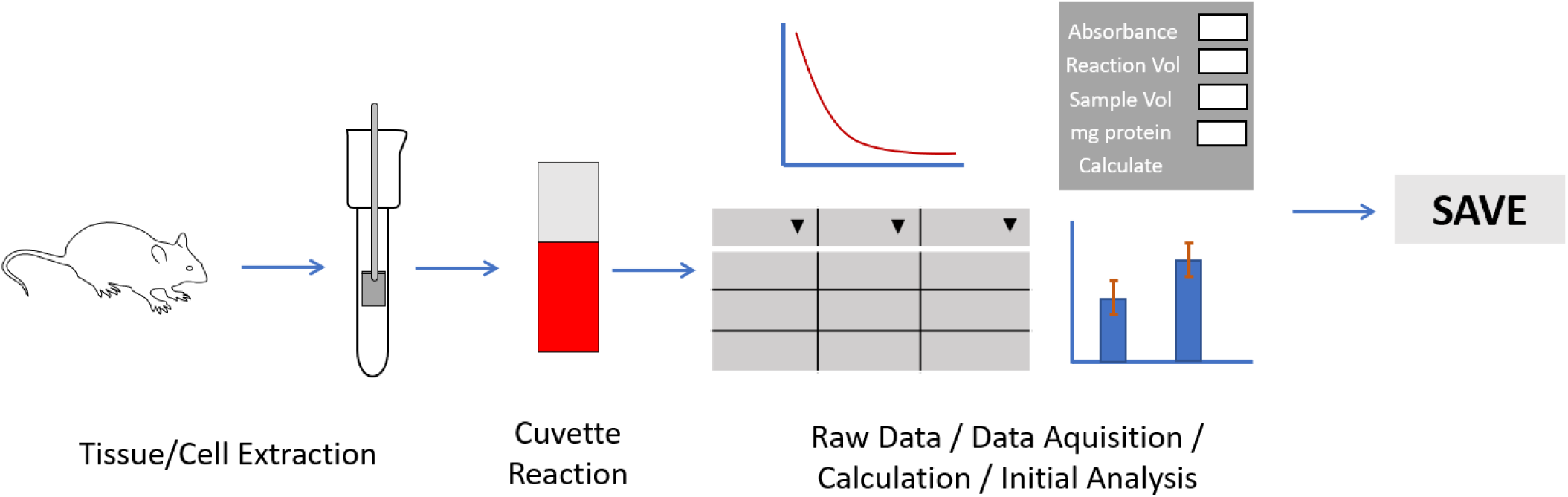

## 1. Introduction

Free radical molecules are highly reactive chemical species that are continuously generated in biological systems. Their excessive accumulation within cells leads to a condition known as oxidative stress, which is characterized by increased oxidation of lipids, proteins, and nucleic acids (DNA) [1]. Cellular free radicals may originate from oxygen, forming reactive oxygen species (ROS), or from nitrogen, forming reactive nitrogen species. These reactive molecules can accumulate due to increased production or impaired removal by antioxidant defense mechanisms [2].

To counteract oxidative stress, cells possess a robust antioxidant system that prevents or repairs oxidative damage. A major component of this system is an enzymatic detoxification pathway that constitutes the first line of defense against ROS ([2]). The principal antioxidant enzymes involved in this process are superoxide dismutase (SOD), catalase (CAT), and glutathione peroxidase (GPx), which act sequentially to neutralize ROS. SOD catalyzes the conversion of superoxide radicals into hydrogen peroxide, while catalase and glutathione peroxidase convert hydrogen peroxide into water, with catalase uniquely producing molecular oxygen as a by-product [3].

Superoxide dismutase exists primarily in two isoforms: a cytosolic copper- and zinc-containing enzyme (CuZnSOD) and a mitochondrial manganese-containing enzyme (MnSOD). Catalase is distributed across several subcellular compartments, including peroxisomes, cytoplasm, and mitochondria, while glutathione peroxidase is localized in the cytosol, mitochondria, and nucleus [2,3]. Due to their central role in maintaining cellular redox homeostasis, numerous experimental approaches have been developed to quantify the activity of these enzymes in biological samples [4].

It is important to emphasize that changes in antioxidant enzyme mRNA or protein expression do not necessarily correspond to changes in enzymatic activity. For example, induction of MnSOD gene expression does not always result in increased enzymatic activity [5]. Therefore, direct quantification of enzyme activity is essential to accurately assess the biological impact of oxidative stress and antioxidant defenses. Such measurements require analytical approaches that demonstrate high sensitivity, accuracy, and reproducibility, as deficiencies in analytical procedures may compromise the interpretation of experimental results.

In this context, computational tools and graphical user interfaces (GUIs) can play an important role in improving analytical workflows and ensuring reliable data analysis. Python is a robust, versatile, and user-friendly programming language that has gained widespread adoption in scientific research due to its accessibility, extensive community support, and broad ecosystem of scientific libraries [6]. In recent years, Python-based tools have been successfully developed for applications in molecular modeling, enzymatic catalysis, cellular data analysis, and protease activity quantification [7–9].

Despite these advances, no freely available, integrated graphical software tool currently exists that is specifically designed to analyze experimental data from antioxidant enzyme activity assays without reliance on spreadsheets or manual calculations. Spreadsheet-based analysis is widely used but is prone to accidental data modification, calculation errors, and inefficiencies when handling multiple samples or experimental conditions. These limitations highlight the need for a standardized, user-friendly computational tool capable of automating antioxidant enzyme activity calculations while maintaining transparency and reproducibility. In this study, we present Redoxyme, a free and open-source Python-based GUI (windows or accessible through web) designed to facilitate and standardize the calculation of superoxide dismutase, catalase, and glutathione peroxidase activities. Redoxyme integrates protein normalization, enzyme-specific calculation routines, data visualization, and data export functionalities within an intuitive windows interface that does not require programming expertise, thereby addressing a critical gap in redox biology data analysis.

## 2. Materials and Methods

Detailed experimental protocols, buffer compositions, reagent concentrations, and incubation conditions for all assays are provided in the Supplementary Methods.

### 2.1 Experimental samples and validation strategy

To validate the accuracy and applicability of Redoxyme, antioxidant enzyme activity data were obtained from rodents (mice and rats) cardiac tissue homogenates routinely used in our laboratory for oxidative stress studies. All animal procedures were conducted in accordance with institutional and national ethical guidelines and were approved by the Institutional Animal Care and Use Committee (IACUC) of the Federal University of Cariri (UFCA), under protocol number 006/2025.

### 2.2 Antioxidant enzyme activity assays

#### 2.2.1 Catalase activity assay

Catalase activity was measured spectrophotometrically by monitoring the decomposition of hydrogen peroxide at 240 nm, following the method originally described elsewher [10]. Cardiac Tissue homogenates were added to a reaction medium containing hydrogen peroxide in phosphate buffer (pH 7.4). The decrease in absorbance was recorded over time, and the linear portion of the decay curve within the first 60 seconds was used for activity calculations. Catalase activity was expressed as units per milligram of protein (U/mg protein).

#### 2.2.2 Superoxide dismutase activity assay

Superoxide dismutase activity was determined using the nitro blue tetrazolium (NBT) reduction inhibition assay, as described by Beauchamp and Fridovich [11]. Tissue homogenates were incubated in a reaction mixture containing riboflavin, methionine, EDTA, and NBT, followed by uniform light exposure. Absorbance was measured at 560 nm. One unit of SOD activity was defined as the amount of enzyme required to inhibit NBT reduction by 50%, and results were normalized to protein content.

#### 2.2.3 Glutathione peroxidase activity assay

Glutathione peroxidase activity was assessed by monitoring the oxidation of NAD(P)H at 340 nm in a coupled enzymatic reaction, as previously described [3,12]. The assay relies on the GPx-mediated reduction of hydrogen peroxide, coupled to glutathione reductase-dependent NAD(P)H oxidation. The decrease in absorbance during the linear phase of the reaction was used to calculate enzyme activity, which was expressed as U/mg protein.

### 2.3 Software implementation and architecture

Redoxyme was developed using Python version 3.9 (Compatible with Python 3.9 and above) and implemented as a graphical user interface using the Tkinter framework. The software integrates several widely used Python libraries, including NumPy for numerical operations, Pandas for data handling, Matplotlib for data visualization, and Openpyxl for Excel file input and output. Current code version 1.0.0. Support email for questions and/or. The permanent link to code/GUI/repository used is https://github.com/hebertyfacundo/redoxyme. Legal Code License MIT License (https://opensource.org/licenses/MIT). The software is distributed as a standalone Windows executable generated using PyInstaller, allowing users to run the application without requiring a Python installation. The full source code is openly available via a public GitHub repository, enabling transparency, inspection, improvements, and future adaptation by the scientific community.

Redoxyme is also a web-based application implemented using the Streamlit framework (https://streamlit.io). The online calculator allows users to perform enzyme activity calculations directly through a browser interface without requiring local installation or specific operating system constraints.

The current stable version of the application is publicly accessible at: https://redoxyme.streamlit.app/

As said above the full source code, version history, and documentation are openly available in a public GitHub repository: https://github.com/hebertyfacundo/redoxyme

### 2.4 Enzyme activity calculation algorithms

Redoxyme implements established mathematical equations for calculating antioxidant enzyme activities based on spectrophotometric measurements. All calculations normalize enzymatic activity to protein concentration, expressed as units per milligram of protein (U/mg protein).

Catalase activity calculations are based on pseudo-first-order kinetics using the natural logarithm of absorbance decay. Glutathione peroxidase activity calculations account for background oxidation by subtracting blank reactions and applying the appropriate molar extinction coefficient for NAD(P)H. Superoxide dismutase activity is calculated based on percent inhibition of NBT reduction relative to control reactions.

All equations implemented in the software are derived from widely accepted biochemical methods and are presented explicitly in the Supplementary Methods, along with worked numerical examples.

### 2.5 Data handling, visualization, and export

Redoxyme allows users to manually enter assay values or import absorbance-versus-time data directly from Excel files. Calculated enzyme activities can be stored within the interface, exported as Excel spreadsheets, and visualized as bar plots displaying mean values with standard deviations. These features enable rapid inspection of results prior to downstream statistical analysis using external software.

## Results

### 3.1 Software workflow and protein normalization

Redoxyme calculates antioxidant enzyme activities by applying standardized equations (please see supplementary data) to individual samples within each experimental condition, reporting enzyme activity as units per milligram of protein (U/mg protein). For each enzyme-specific calculator, users manually enter experimental parameters including absorbance values, reaction volume, sample volume, dilution factor (when applicable), and protein concentration.

Because enzyme activity is intrinsically dependent on protein content, Redoxyme (windows GUI only at this point) incorporates an integrated protein quantification module accessible through the “Protein” button within each enzyme calculator. This module allows users to generate a protein standard curve by entering absorbance values of known protein concentrations. Sample absorbance values are then interpolated automatically to calculate protein concentration, which is subsequently used to normalize enzyme activity values. An example of the protein concentration calculator embedded in Redoxyme is shown in Figure 1. Catalase, glutathione peroxidase or superoxide dismutase activity calculated can be accessed from Redoxyme main window (Figure 2).

**Figure 1:**
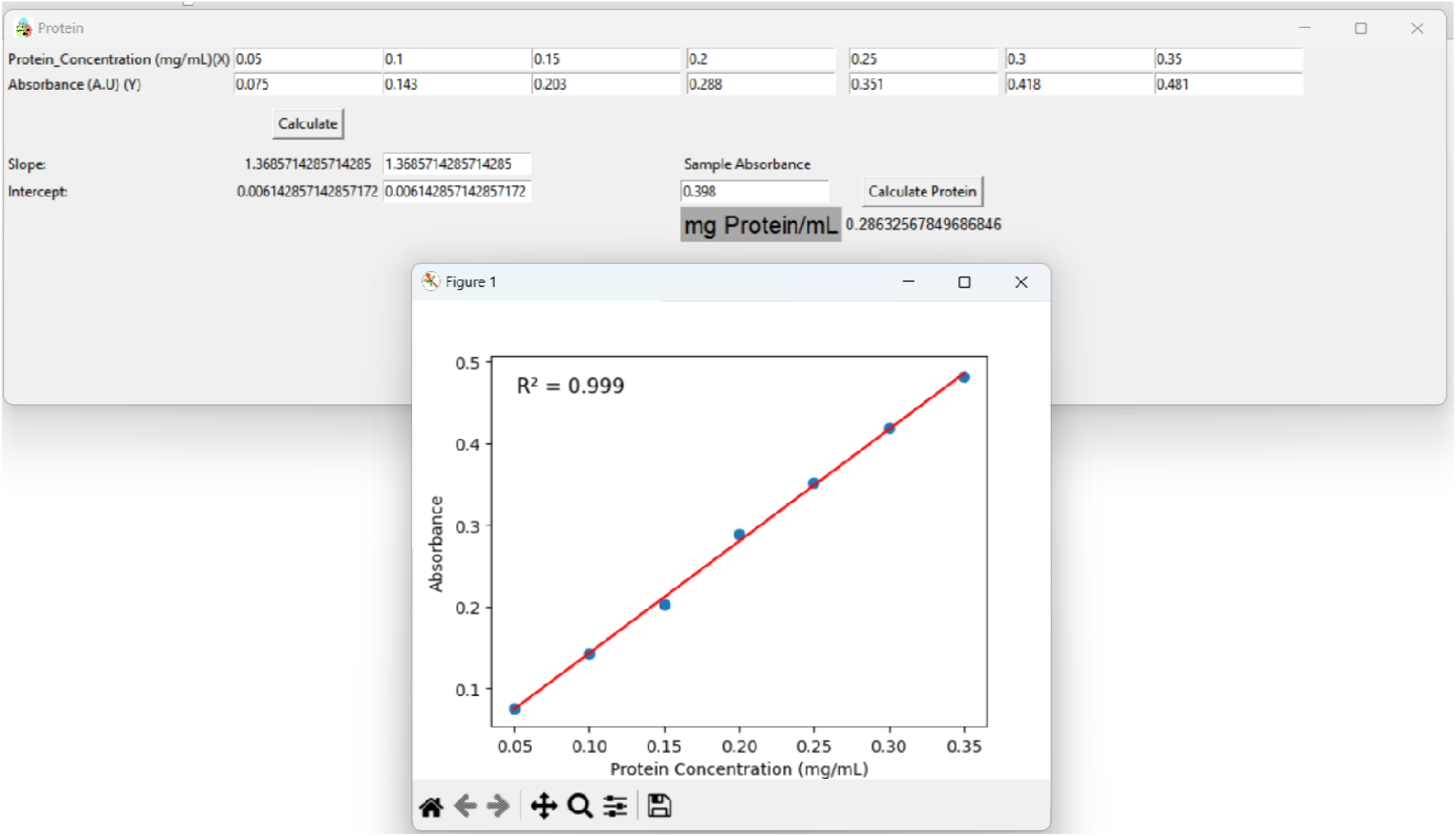
Main window of Redoxyme program. The buttons at the bottom will link to catalase, glutathione peroxidase or superoxide dismutase activity calculator, respectively. The main window of the program has a figure of the reactions catalyzed by each enzyme.

**Figure 2:**
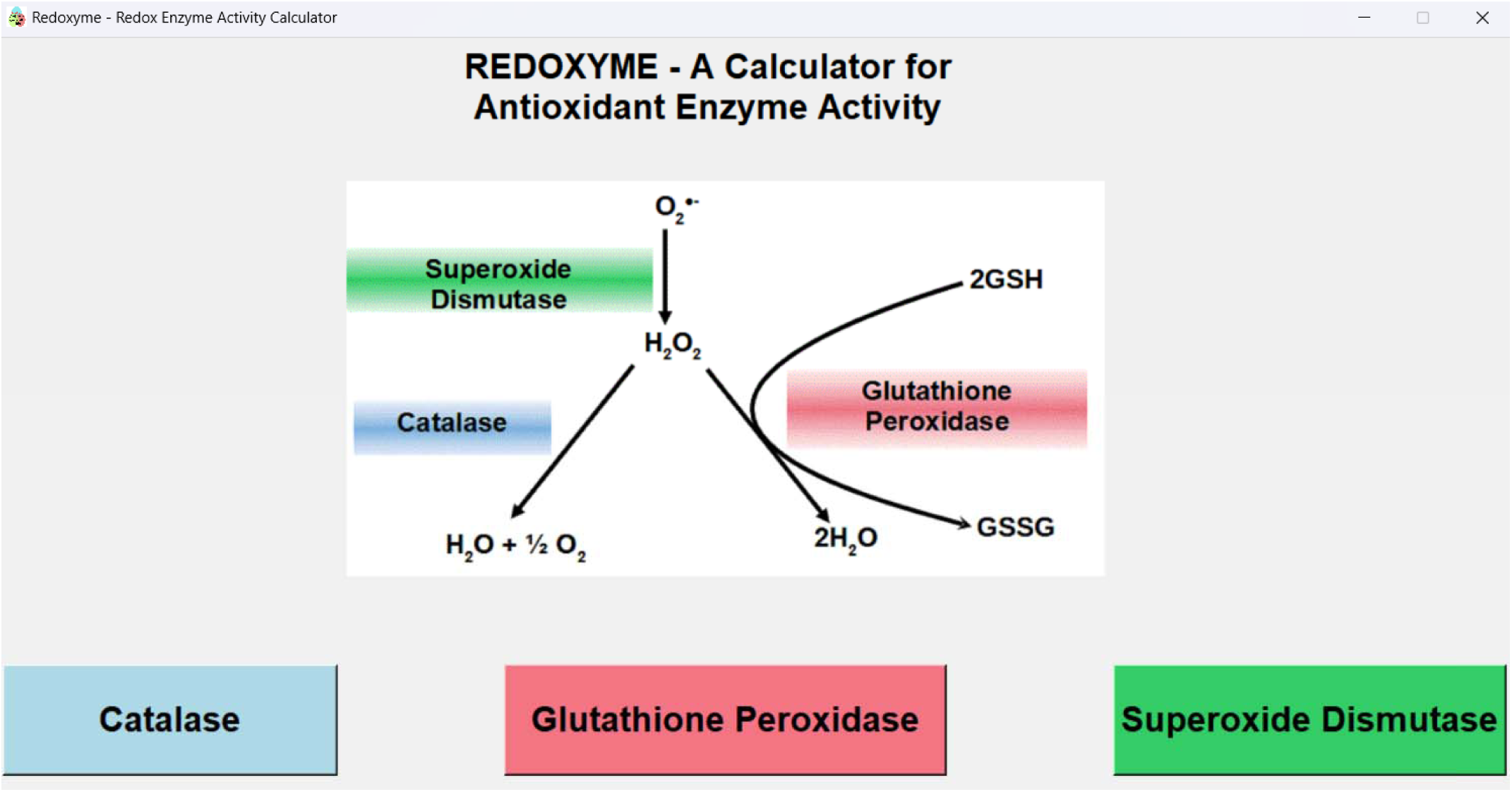
Protein concentration calculator integrated into Redoxyme. The interface generates a standard curve from user-defined protein standards and applies it to sample absorbance values, providing protein normalization directly within the software for subsequent enzyme activity calculations.

### 3.2 Shared features of time-dependent enzyme activity calculators

Catalase and glutathione peroxidase (GPx) activities are determined from time-dependent spectrophotometric assays. In both cases, absorbance values at 0 and 60 seconds are required for activity calculations, as described in the Materials and Methods section. These values may be entered manually. Importantly, redoxyme (windows GUI only) includes a dedicated “Graph XY Excel” function that allows users to visualize absorbance as a function of time after importing Excel data (it must be “time” in column A and “absorbance” in column B). This feature generates a line plot using the Matplotlib library and enables users to identify absorbance values at specific time points by hovering over the curve (Figures 3 and 4 inserts). Users may also zoom, customize, and export the generated plots in standard image formats (e.g., PNG or TIFF). This visualization feature is available for catalase, GPx, and superoxide dismutase (SOD) calculators, although the SOD assay does not rely on time-scan data. During Excel import, Redoxyme may detect values formatted as text rather than numeric values. In such cases, users must convert text to numeric format prior to import, for example by using the “Text to Columns” function in Microsoft Excel.

**Figure 3:**
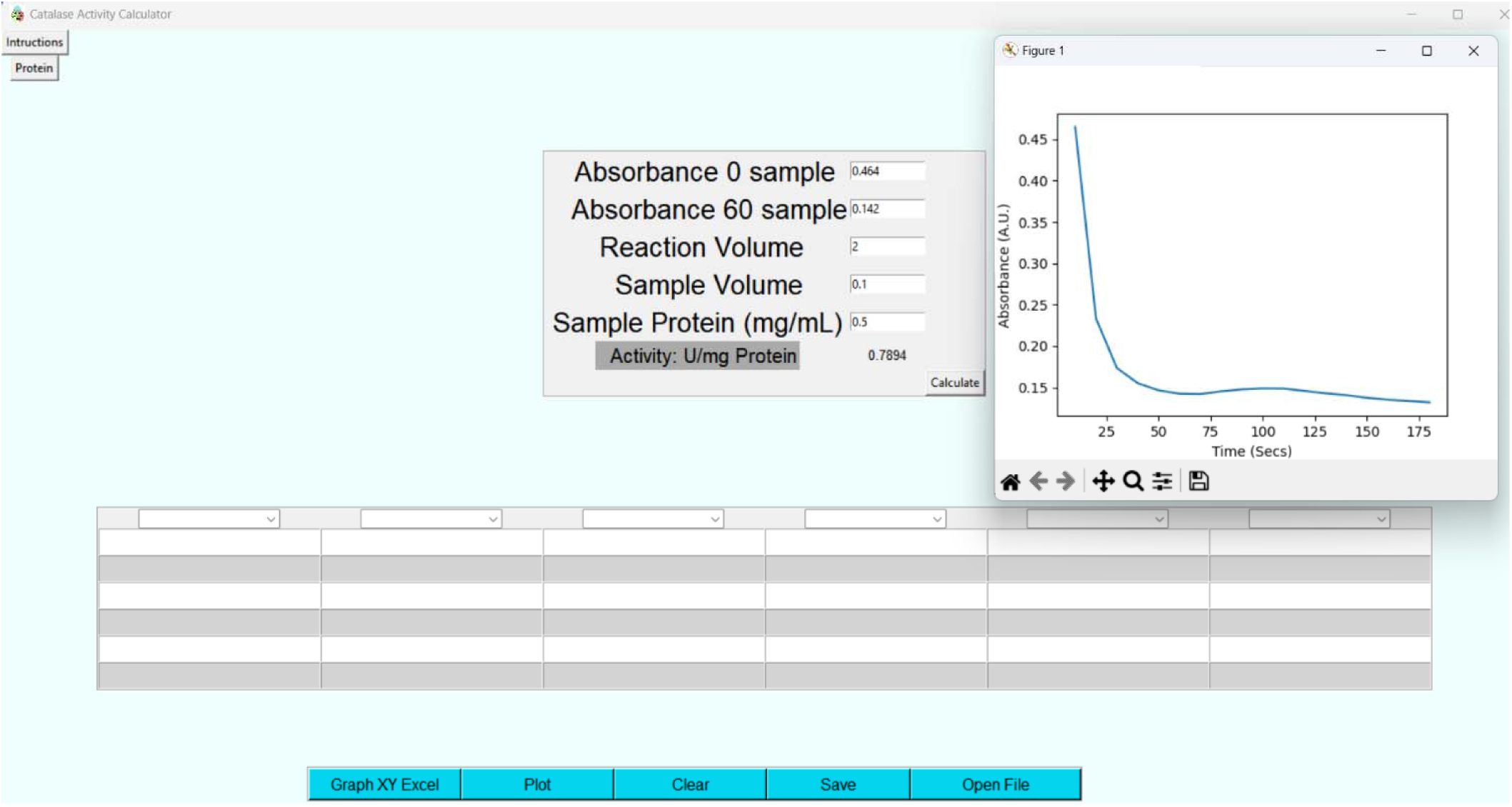
Illustration of the catalase activity calculator window. Graphical user interface of the catalase activity calculator implemented in Redoxyme. The module allows manual input of absorbance values at 0 and 60 seconds, reaction volume, sample volume, and protein concentration, and automatically computes catalase activity normalized to protein content (U/mg protein) using a predefined kinetic equation. A time-dependent absorbance plot generated from imported spectrophotometric data is displayed to assist users in visual inspection and extraction of absorbance values. Calculated activities can be stored in an integrated table (which can be saved in excel format), exported, or visualized in a plot, enabling standardized and reproducible analytical data processing without the use of external spreadsheets.

**Figure 4:**
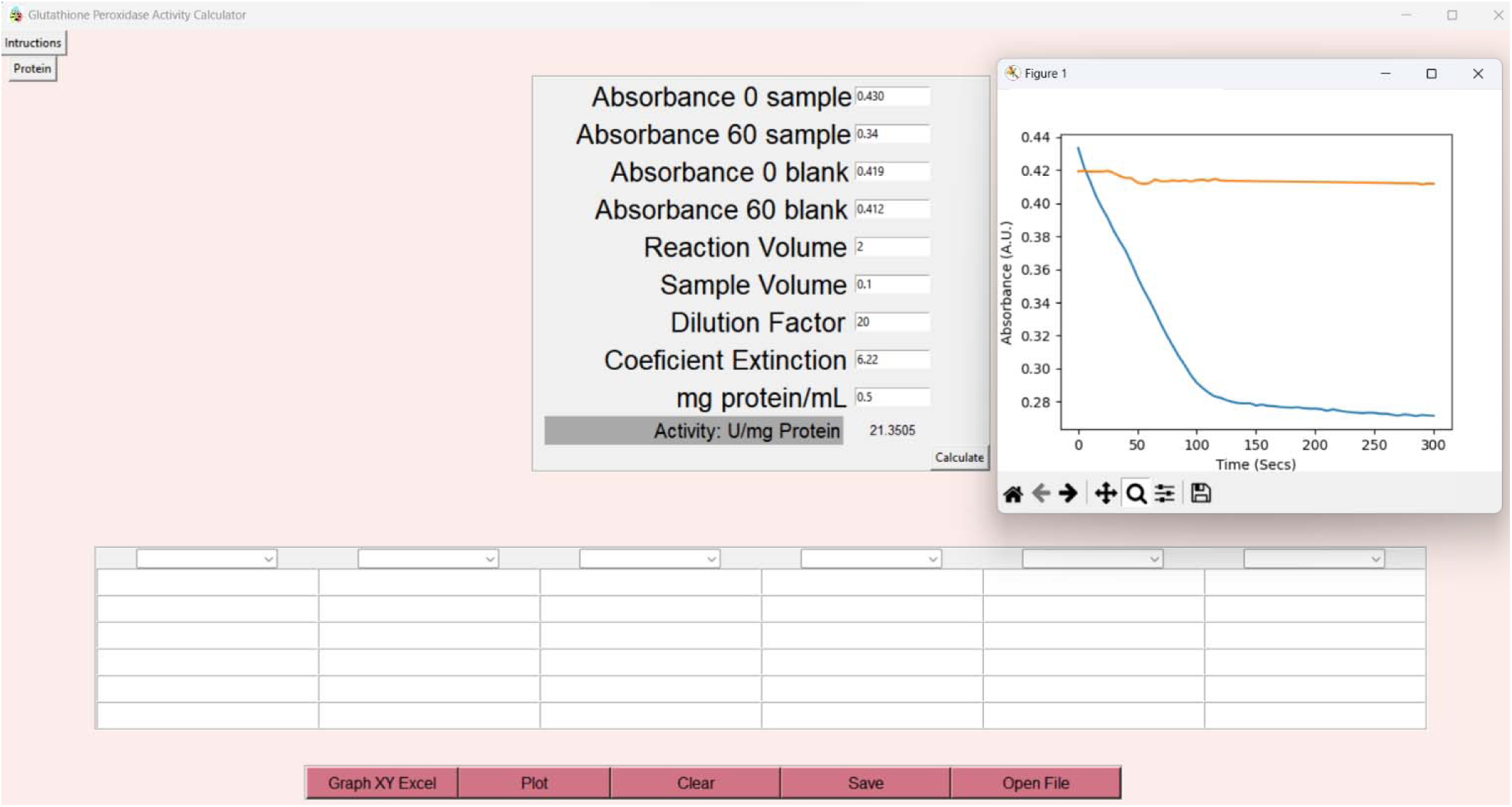
Illustration of the glutathione peroxidase activity calculator window. Graphical user interface of the Glutathione peroxidase activity calculator implemented in Redoxyme. The module allows manual input of absorbance values at 0 and 60 seconds, reaction volume, sample volume, and protein concentration, and automatically computes glutathione peroxidase activity normalized to protein content (U/mg protein) using a predefined kinetic equation. A time-dependent absorbance plot generated from imported spectrophotometric data is displayed to assist users in visual inspection and extraction of absorbance values. Calculated activities can be stored in an integrated table (which can be saved in excel format), exported, or visualized in a plot, enabling standardized and reproducible analytical data processing without the use of external spreadsheets.

### 3.3 Catalase activity calculation and output

The catalase activity calculator is accessed by selecting the “Catalase” option in the main window (Figure 2), which opens a dedicated interface (Figure 3). Users enter absorbance values at 0 and 60 seconds, reaction volume, sample volume, and protein concentration. Upon clicking the “Calculate” button, Redoxyme computes catalase activity and displays the result as U/mg protein.

Catalase activity is calculated using a pseudo–first-order kinetic equation that converts hydrogen peroxide consumption into enzymatic units, as described by Aebi[10]. The equations implemented in Redoxyme are provided in the Supplementary Methods. Calculated values (windows GUI) are automatically copied to the system clipboard, allowing users to paste results directly into external software such as GraphPad Prism or Excel. Alternatively, results can be entered into the internal results table displayed in the catalase calculator window. The user may build graphs from the data for inspection of publication.

### 3.4 Glutathione peroxidase activity calculation and output

The glutathione peroxidase calculator is accessed via the “Glutathione Peroxidase” button in the main interface, opening a dedicated input window (Figure 4). Required inputs include absorbance values at 0 and 60 seconds for both blank and sample reactions, reaction volume, sample volume, dilution factor, and protein concentration. After calculation, GPx activity is displayed as U/mg protein.

GPx activity is calculated based on the rate of NAD(P)H oxidation at 340 nm, using the appropriate molar extinction coefficient. Redoxyme automatically applies blank correction and dilution factor adjustments. As with catalase, calculated GPx activity values are copied to the clipboard and can also be stored in the internal results table. Full equations and worked numerical examples are provided in the Supplementary Methods.

### 3.5 Superoxide dismutase activity calculation and output

The superoxide dismutase activity calculator is accessed through the “Superoxide Dismutase” button in the main window (Figure 2), opening a dedicated input interface (Figure 5). Users enter absorbance values for blank and sample reactions, reaction volume, sample volume, dilution factor, and protein concentration. Redoxyme calculates SOD activity based on the percent inhibition of nitro blue tetrazolium reduction, with one unit of SOD defined as the amount of enzyme required to produce 50% inhibition. Calculated SOD activity values are displayed as U/mg protein and may be copied to the clipboard or stored in the internal results table (windows GUI). The equations implemented in the SOD calculator are detailed in the Supplementary Methods.

**Figure 5:**
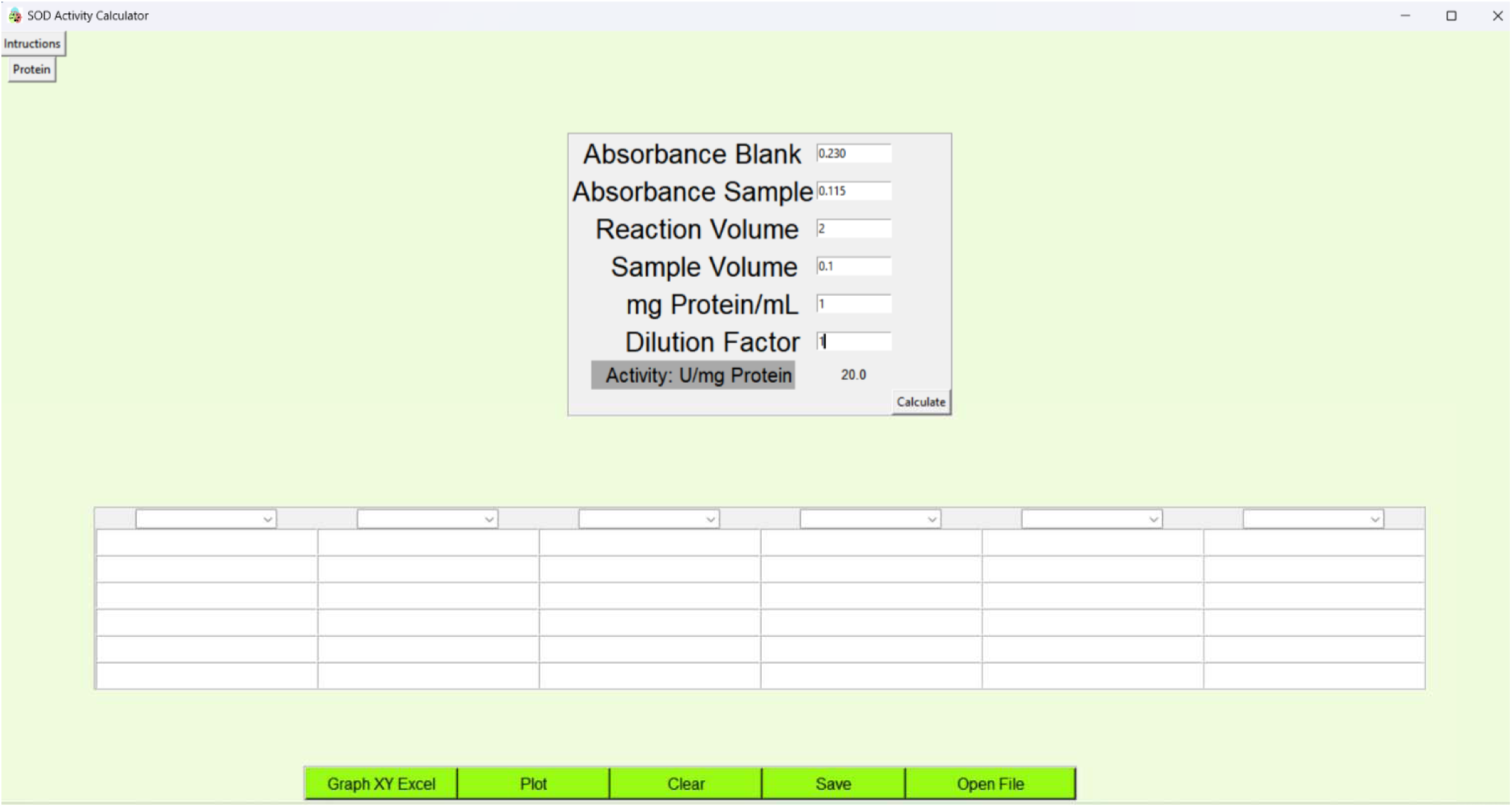
Illustration of the superoxide dismutase activity calculator window. Graphical user interface of the Glutathione peroxidase activity calculator implemented in Redoxyme. The module allows manual input of absorbance values at 0 and 60 seconds, reaction volume, sample volume, and protein concentration, and automatically computes superoxide dismutase activity normalized to protein content (U/mg protein) using a predefined kinetic equation. Calculated activities can be stored in an integrated table (which can be saved in excel format), exported, or visualized in a plot, enabling standardized and reproducible analytical data processing without the use of external spreadsheets.

### 3.6 Data storage, export, and visualization

All enzyme activity calculators windows within Windows graphical user interface of Redoxyme include an internal table for organizing calculated results across multiple samples and experimental conditions. Users may assign custom labels to each column or using using dropdown menus (e.g., control, treated). Data tables can be saved at any time as Excel files, which are automatically named according to enzyme type and timestamp and stored in the same directory as the executable file.

### 3.7 Test to experimental data

Redoxyme was applied to data obtained from mice and rat heart samples (Figure 6A, B and C). The software successfully processed all datasets, producing enzyme activity values consistent with manual calculations and standard analytical procedures. These results demonstrate the applicability of Redoxyme to both biologically complex samples and controlled enzymatic systems.

**Figure 6.**
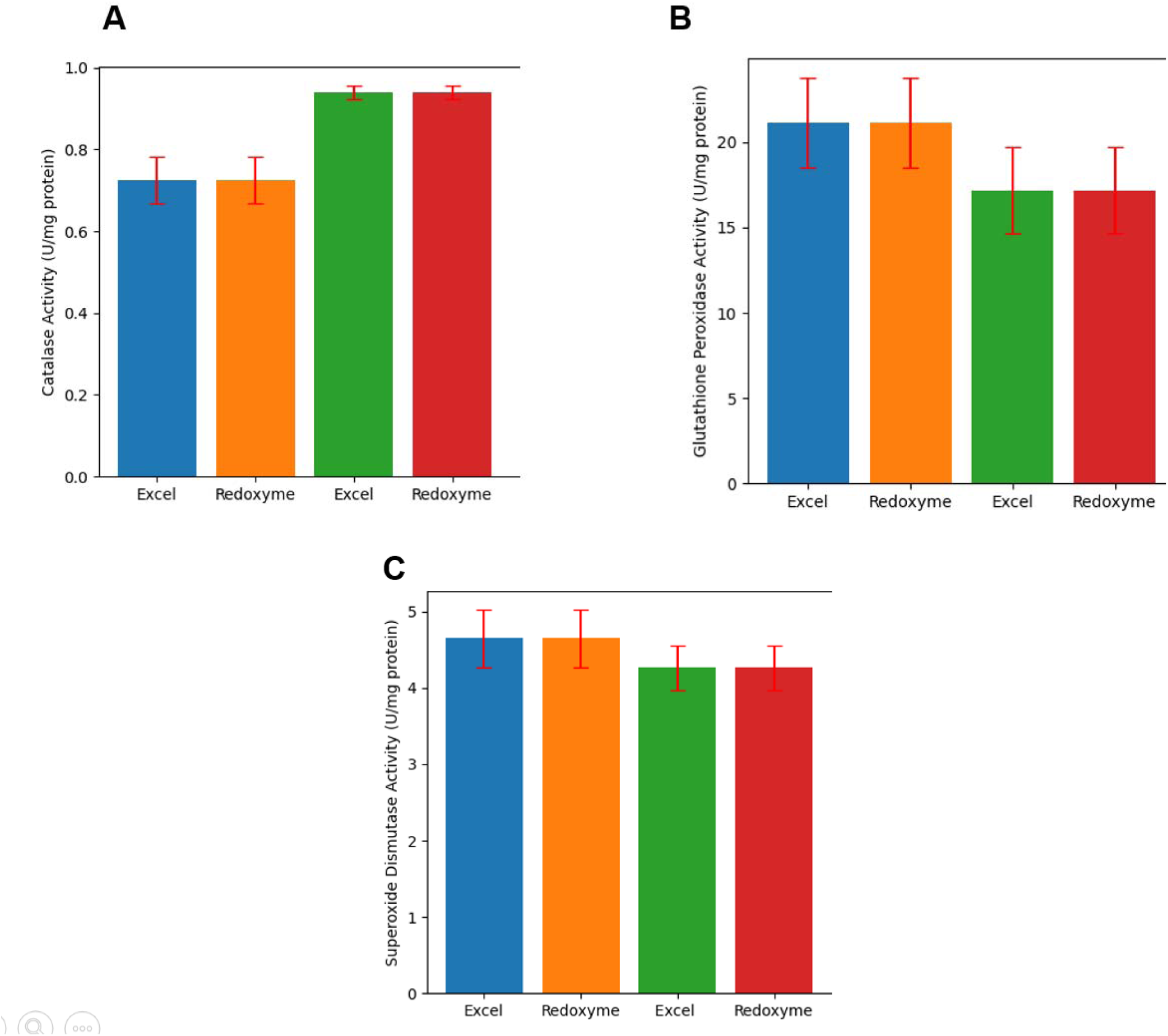
Validation of Redoxyme enzyme activity calculations using experimental heart homogenates. Comparison of antioxidant enzyme activities calculated using conventional spreadsheet-based analysis (Excel) and the Redoxyme software in heart homogenates from mice and rats. (A) Catalase activity, (B) glutathione peroxidase activity, and (C) superoxide dismutase activity. In each panel, the first and second bars correspond to mouse samples analyzed using Excel and Redoxyme, respectively, while the third and fourth bars correspond to rat samples analyzed using Excel and Redoxyme. Enzyme activities are expressed as Units per milligram of protein (U/mg protein). Data are presented as mean ± standard deviation (n = 3 independent samples per group).

## 4. Discussion

In this study, we present Redoxyme, a lightweight, open-source Python-based graphical user interface designed to facilitate the analysis of antioxidant enzyme activities, specifically glutathione peroxidase, catalase, and superoxide dismutase. The primary objective of the software is to streamline enzymatic activity calculations while minimizing reliance on spreadsheets and manual data handling. Redoxyme provides a simple and stable workflow that allows users to visualize time-scan data, extract numerical values, calculate enzyme activities normalized to protein content (which may be also calculated inside the program), store results in structured tables, and generate graphical outputs. An overview of the software workflow is presented in Figure 7.

**Figure 7.**
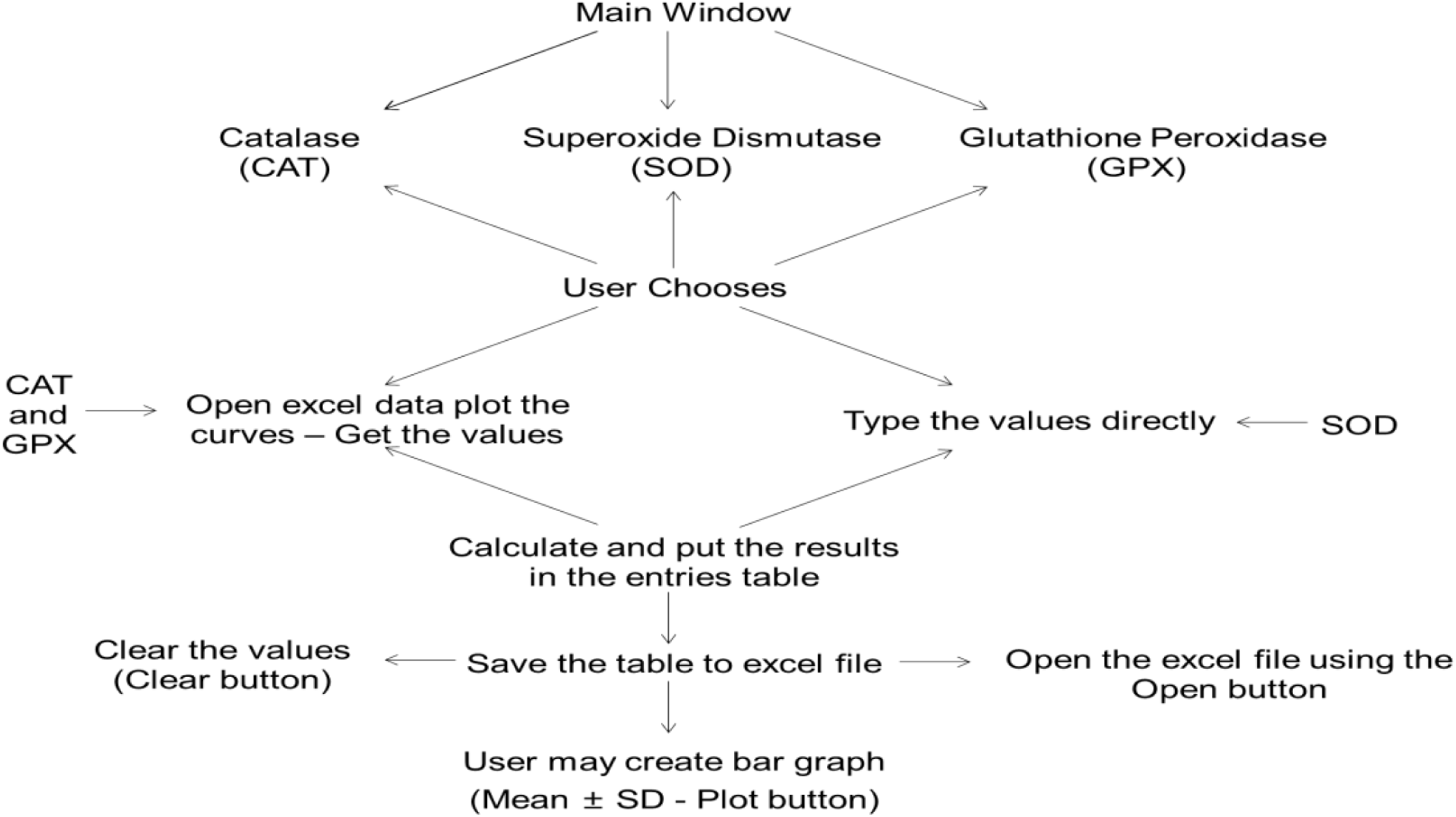
Workflow of Redoxyme. Schematic representation of the Redoxyme workflow from user interaction to data output. From the main window, users select the enzyme activity module (catalase, superoxide dismutase, or glutathione peroxidase). Depending on the assay, absorbance values are either entered directly or extracted (by the user) from time-scan data imported from Excel files. Redoxyme automatically performs enzyme activity calculations and stores results in an integrated table, which can be saved as an Excel file, reopened, or cleared. The software also enables graphical visualization of results as bar plots displaying mean ± standard deviation.

Oxidative stress plays a critical role in the pathophysiology of numerous human diseases and aging processes. Reactive oxygen species are generated physiologically and participate in essential signaling pathways, but their excessive accumulation can result in cellular damage [13]. Enzymatic antioxidants such as catalase, superoxide dismutase, and glutathione peroxidase constitute a primary defense system against oxidative stress and are routinely quantified through activity-based assays [4,14]. Despite the widespread use of these assays, the computational analysis of antioxidant enzyme activity data is still predominantly performed using spreadsheets, which increases the risk of calculation errors and inconsistencies. The enzymatic activity assays implemented in Redoxyme [10–12] were selected based on their extensive adoption in the literature. While alternative protocols exist (for instance, in gel activity protocols) for each enzyme [3], the reference methodologies for catalase, superoxide dismutase, and glutathione peroxidase rank among the most highly cited protocols in redox biology (Supplementary Table S1), supporting their status as canonical assays and ensuring compatibility with the majority of published workflows and facilitating direct comparison with existing literature.

A major contribution of Redoxyme is the standardization of enzyme activity calculations through the implementation of validated equations within a controlled computational environment. Spreadsheet-based analyses are known to be vulnerable to accidental modification, copy-paste errors, and formula inconsistencies, particularly when datasets increase in size or complexity. By embedding enzyme-specific calculation logic directly into the software, Redoxyme reduces these risks and promotes reproducible data analysis. This feature is particularly relevant for laboratories with multiple users or for training environments involving undergraduate and graduate students. Importantly, the publicly accessible web-based platform (redoxyme.streamlit.app) represents an important step toward improving accessibility and standardization of enzymatic activity calculations. By integrating open-source distribution through GitHub with a browser-based interface, Redoxyme reduces technical barriers associated with installation and operating system dependencies. Furthermore, open version control enhances transparency, allowing users to audit computational routines and reproduce analyses across independent laboratories

Importantly, Redoxyme was not developed to replace spreadsheets entirely or to diminish their educational value. Spreadsheet software remains a valuable tool for data organization, visualization, and teaching fundamental concepts. However, for routine and repetitive enzymatic activity calculations, dedicated software solutions can substantially improve efficiency and reliability. Redoxyme eliminates repetitive data transfer steps, minimizes manual intervention, and reduces the likelihood of accidental data corruption, thereby improving overall analytical robustness.

The graphical user interface was intentionally designed to prioritize usability and low cognitive load, enabling researchers without programming expertise to perform complex calculations efficiently. Well-designed graphical interfaces are known to improve task completion speed and reduce user error. While proficiency in programming languages such as Python, R, or C++ offers clear advantages for data analysis [6], many experimental researchers rely on GUIs to visualize, analyze, and archive their data. Redoxyme directly addresses this need within the field of redox biology by providing an accessible tool for antioxidant enzyme activity analysis.

Performance analysis demonstrated that Redoxyme executes enzyme activity calculations with minimal computational overhead. Average execution times were approximately 0.00501seconds for catalase calculations, 0.002001 seconds for glutathione peroxidase calculations, and 0.00399 seconds for superoxide dismutase calculations, demonstrating the lightweight computational footprint and rapid responsiveness of the software. These execution times indicate that the software’s responsiveness is limited primarily by user interaction rather than computational constraints, which is advantageous for routine laboratory use and high-throughput experimental workflows.

At present, Redoxyme is distributed as a standalone executable for the Windows operating system. While this represents a limitation, it reflects the availability of development and testing resources at the time of implementation. The web-based Streamlit Application implementation provides platform-independent access via web browsers (for enzymatic activity calculation only). While the web-based Streamlit application provides a practical and readily accessible platform for rapid enzyme activity calculations, the locally executed Windows version retains full analytical functionality, including integrated protein normalization, graphical visualization of experimental data, and expanded workflow tools designed for comprehensive laboratory data processing. An inherent limitation of Redoxyme is that its algorithms are based on defined enzymatic activity assays [10–12], requiring users to adopt the corresponding experimental protocols. Alternative methodologies may therefore fall outside the current scope of the software. Nevertheless, the open-source architecture of Redoxyme allows future adaptation of the computational models to accommodate additional assay formats, enabling methodological expansion without compromising standardization. Importantly, the software was developed entirely using open-source Python libraries, and the source code is publicly available, allowing future adaptation to other operating systems such as macOS or Linux. This transparency supports long-term sustainability and community-driven development.

## 5. Conclusion

In conclusion, Redoxyme provides a practical and reproducible solution for the calculation of antioxidant enzyme activities. By integrating standardized calculations, protein normalization, data visualization, and export functionality within a single graphical interface, Redoxyme enhances analytical consistency while reducing user-dependent variability. The software is particularly well suited for academic laboratories, teaching environments, and clinical research settings where simplicity, speed, and reliability are essential. Future development efforts may focus on cross-platform compatibility and expanded support for additional redox-related assays.

## Data Availability

The windows graphical user interfac and the open-source codes (for windows and streamlit web application) are fully available at https://github.com/hebertyfacundo/redoxyme. Raw files are made available upon reasonable request.

## Author Contributions

G.C.F.S - formal analysis, investigation, methodology; A.L.N.V - investigation, methodology H.T.F. – supervision, writing, original draft, software writing - review & editing. All authors read and approved the final manuscript.

---

**Table S1.** Scopus citation metrics for reference antioxidant enzyme assays. Searched in 01 31^st^ 2026.

| Enzyme | Reference | Scopus Citation Count* |
| --- | --- | --- |
| Catalase | Aebi 1984 | 23492 |
| Superoxide Dismutase | Beauchamp & Fridovich 1971 | 11739 |
| Glutathione Peroxidase | Paglia 1967 | 10791 |

## Supplementary Methods

### S1. Tissue Sample Preparation

The heart tissue was minced into small pieces using scissors, washed twice with ice cold PBS and homogenized with a precooled glass potter (40 strokes) in ice-cold buffer composed of 300mM sucrose, 10mM K^+^Hepes buffer, pH 7.2, 1mM K^+^EGTA, and BSA 1 mg/mL. Nuclei and cellular residues were pelleted by centrifugation at 1200×g for 5 min. The supernatant was recentrifuged at 9300 g for 10 min. The resultant supernatant (cytosolic fraction) was stored immediately at −80°C and used catalase, superoxide dismutase or glutathione peroxidase activity.

### S2. Detailed antioxidant enzyme activity assay protocols

#### S2.1 Catalase activity assay

Catalase activity was determined by monitoring the decomposition of hydrogen peroxide (H₂O₂) at 240 nm using a spectrophotometric method originally described by Aebi (Aebi H., Catalase in vitro, Methods Enzymol, 105:121–126, 1984) Cited by 23492. Cardiac tissue homogenates were added to a reaction medium containing hydrogen peroxide prepared in phosphate buffer (100 mM, pH 7.4). The final concentration of H₂O₂ in the reaction mixture was 50 mM. Absorbance changes were recorded for 2–3 minutes at 240 nm using a spectrophotometer. The initial absorbance was approximately 0.5 at the start of the reaction. Catalase activity was calculated using the linear portion of the decay curve during the first 60 seconds. Enzyme activity was expressed as units per milligram of protein (U/mg protein).

#### S2.2 Superoxide dismutase activity assay

Superoxide dismutase (SOD) activity was measured using the nitro blue tetrazolium (NBT) reduction inhibition assay described by Beauchamp and Fridovich (Beauchamp C., Fridovich I., Anal Biochem, 44:276–287, 1971) Cited by 11739. Clear tissue homogenate supernatants were added to a reaction mixture containing 0.1 mM EDTA, 13 mM L-methionine, and 75 µM NBT in potassium phosphate buffer (pH 7.8). The reaction was initiated by the addition of 2 µM riboflavin, followed by uniform exposure to non-filtered white light for 5 minutes. The formation of reduced NBT was measured at 560 nm. One unit of SOD activity was defined as the amount of enzyme required to inhibit NBT reduction by 50%. Results were normalized to protein concentration and expressed as U/mg protein. For mitochondrial MnSOD activity determination, samples were subjected to three freeze–thaw cycles in hypotonic buffer (25 mM K₂PO₄, pH 7.2; 5 mM MgCl₂). Samples were then incubated with 5 mM potassium cyanide (KCN) for 20 minutes to inhibit cytosolic CuZnSOD activity prior to analysis.

#### S2.3 Glutathione peroxidase activity assay

Glutathione peroxidase (GPx) activity was measured by monitoring NAD(P)H oxidation at 340 nm in a coupled enzymatic reaction Paglia DE, Valentine WN. Studies on the quantitative and qualitative characterization of erythrocyte glutathione peroxidase. J Lab Clin Med. 1967 Jul;70(1):158-69. PMID: 6066618. (Cited by 10791 scopus) And Weydert C.J., Cullen J.J., Nat Protoc, 5:51–66, 2010) Cited. 1,272 ScopusP The reaction buffer consisted of 50 mM potassium phosphate, 0.5 mM EDTA, 1 mM reduced glutathione (GSH), 1.5 mM sodium azide, and 0.2 U/mL glutathione reductase (Sigma-Aldrich), adjusted to pH 7.0. NADH was used at a final concentration of 40 mM. Cell or tissue homogenates (0.3–1 mg protein) were incubated in the reaction mixture for 5 minutes prior to reaction initiation. The reaction was initiated by adding hydrogen peroxide or organic peroxide substrates (e.g., cumene hydroperoxide or tert-butyl hydroperoxide). Absorbance at 340 nm was monitored for 5 minutes, and the linear decay between 0 and 60 seconds was used to calculate enzyme activity. GPx activity was expressed as U/mg protein using a molar extinction coefficient of 6.22 mM⁻¹·cm⁻¹ for NAD(P)H.

### S3. Protein quantification method

Protein concentration was determined spectrophotometrically using a standard curve generated from known protein concentrations. Absorbance values of protein standards were measured, and linear regression analysis was used to calculate the slope and intercept of the standard curve. Sample absorbance values were then interpolated using the regression equation to determine protein concentration. Redoxyme incorporates a dedicated protein quantification module that allows users to input standard absorbance values and concentrations to generate a calibration curve. Sample absorbance values are subsequently entered, and protein concentration is calculated automatically. These values are used to normalize antioxidant enzyme activities, expressed as units per milligram of protein (U/mg protein).

### S3. Enzyme activity calculation equations and worked examples

#### S3.1 Catalase activity calculation

Catalase activity was calculated based on pseudo-first-order kinetics using the following equation:

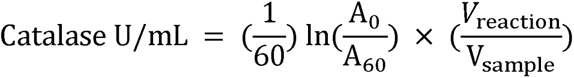

Catalase activity normalized to protein concentration was calculated as:

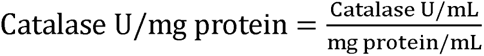

**Example:**

A₀ = 0.464

A₆₀ = 0.142

Reaction volume = 2.0 mL

Sample volume = 0.1 mL

Protein concentration = 0.5 mg/mL

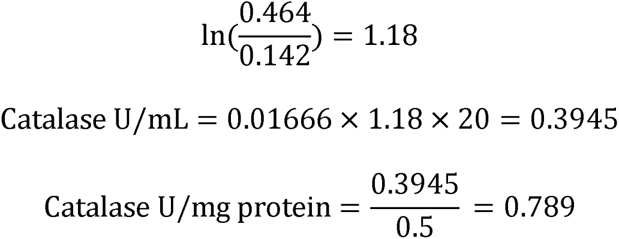

#### S3.2 Glutathione peroxidase activity calculation

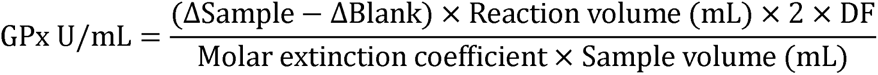

Where:

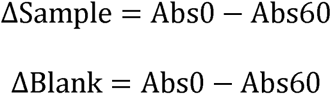

GPx activity was normalized to protein concentration:

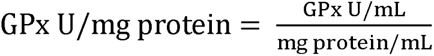

**Worked example:**

Abs Sample₀ = 0.430 Abs

Sample₆₀ = 0.340

Abs Blank₀ = 0.419

Abs Blank₆₀ = 0.412

Reaction volume = 2.0 mL

Sample volume = 0.1 mL

Dilution factor = 20

Protein concentration = 0.5 mg/mL

Final calculated GPx activity = **21.35 U/mg protein**.

**S3.3 Superoxide dismutase activity calculation**

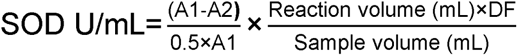

Where:

- A₁ = absorbance of blank reaction
- A₂ = absorbance of sample reaction
- 0.5 represents 50% inhibition of NBT reduction

**Worked example:**

Abs Blank = 0.230

Abs Sample = 0.115

Reaction volume = 2.0 mL

Sample volume = 0.1 mL

Dilution factor = 1

Protein concentration = 1 mg/mL

Final calculated SOD activity = **20 U/mg protein**.

### S4. Software installation and execution (Windows)

Redoxyme is distributed as a standalone Windows executable. Users must download all compressed archive files from the GitHub repository (https://github.com/hebertyfacundo/redoxyme) and extract them using a decompression utility such as 7-Zip. After extraction, the executable file (Redoxyme.exe) can be launched directly without requiring Python installation. The icon file must remain in the same directory as the executable to ensure proper file saving and graphical display. Output files generated by the software are saved in the same directory by default.

### S5. Runtime benchmarking and performance assessment

The execution time of individual enzyme activity calculations was assessed using Python timing functions from the time and datetime modules. Execution time was measured as the elapsed time between the start and end of calculation functions.

## References

[1] H.J. Forman, H. Zhang, Targeting oxidative stress in disease: promise and limitations of antioxidant therapy, Nat. Rev. Drug Discov. 20 (2021) 689–709. 10.1038/S41573-021-00233-1.

[2] H. Sies, Oxidative stress: A concept in redox biology and medicine, Redox Biol. 4 (2015) 180–183. 10.1016/j.redox.2015.01.002.

[3] C.J. Weydert, J.J. Cullen, Measurement of superoxide dismutase, catalase and glutathione peroxidase in cultured cells and tissue, Nat. Protoc. 5 (2010) 51–66. 10.1038/NPROT.2009.197.

[4] C. Vives-Bauza, A. Starkov, E. Garcia-Arumi, Measurements of the Antioxidant Enzyme Activities of Superoxide Dismutase, Catalase, and Glutathione Peroxidase, Methods Cell Biol. 80 (2007) 379–393. 10.1016/S0091-679X(06)80019-1.

[5] M.J. Czaja, M.L. Schilsky, Y. Xu, P. Schmiedeberg, A. Compton, L. Ridnour, L.W. Oberley, Induction of MnSOD gene expression in a hepatic model of TNF-alpha toxicity does not result in increased protein, Am. J. Physiol. 266 (1994). 10.1152/AJPGI.1994.266.4.G737.

[6] J.M. Perkel, Programming: Pick up Python, Nature 518 (2015) 125–126. 10.1038/518125A.

[7] F. Zahariev, N. De Silva, M.S. Gordon, T.L. Windus, M. Dick-Perez, ParFit: A Python-Based Object-Oriented Program for Fitting Molecular Mechanics Parameters to ab Initio Data, J. Chem. Inf. Model. 57 (2017) 391–396. 10.1021/ACS.JCIM.6B00654.

[8] A.P. Soleimany, C. Martin-Alonso, M. Anahtar, C.S. Wang, S.N. Bhatia, Protease Activity Analysis: A Toolkit for Analyzing Enzyme Activity Data, ACS Omega 7 (2022) 24292–24301. 10.1021/ACSOMEGA.2C01559.

[9] M.D. Olp, K.S. Kalous, B.C. Smith, ICEKAT: an interactive online tool for calculating initial rates from continuous enzyme kinetic traces, BMC Bioinformatics 21 (2020). 10.1186/S12859-020-3513-Y.

[10] H. Aebi, Catalase in vitro., Methods Enzymol. 105 (1984) 121–126. http://www.ncbi.nlm.nih.gov/pubmed/6727660.

[11] C. Beauchamp, I. Fridovich, Superoxide dismutase: improved assays and an assay applicable to acrylamide gels., Anal. Biochem. 44 (1971) 276–287. http://www.ncbi.nlm.nih.gov/pubmed/4943714.

[12] D.E. Paglia, W.N. Valentine, Studies on the quantitative and qualitative characterization of erythrocyte glutathione peroxidase - PubMed, J Lab Clin Med. 70 (1967) 158–169. https://pubmed.ncbi.nlm.nih.gov/6066618/ (accessed January 31, 2026).

[13] Y.A. Hajam, R. Rani, S.Y. Ganie, T.A. Sheikh, D. Javaid, S.S. Qadri, S. Pramodh, A. Alsulimani, M.F. Alkhanani, S. Harakeh, A. Hussain, S. Haque, M.S. Reshi, Oxidative Stress in Human Pathology and Aging: Molecular Mechanisms and Perspectives, Cells 11 (2022). 10.3390/CELLS11030552.

[14] C.E.B. David, A.M.B. Lucas, M.T.S. Araújo, B.N. Coelho, J.B.S. Neto, B.R.C. Portela, A.L.N. Varela, A.J. Kowaltowski, H.T. Facundo, Calorie restriction attenuates hypertrophy-induced redox imbalance and mitochondrial ATP-sensitive K+ channel repression, J. Nutr. Biochem. 62 (2018) 87–94. 10.1016/j.jnutbio.2018.08.008.

